# False Discovery Rates: A New Deal

**DOI:** 10.1101/038216

**Authors:** Matthew Stephens

## Abstract

We introduce a new Empirical Bayes approach for large-scale hypothesis testing, including estimating False Discovery Rates (FDRs), and effect sizes. This approach has two key differences from existing approaches to FDR analysis. First, it assumes that the distribution of the actual (unobserved) effects is unimodal, with a mode at 0. This “unimodal assumption” (UA), although natural in many contexts, is not usually incorporated into standard FDR analysis, and we demonstrate how incorporating it brings many benefits. Specifically, the UA facilitates efficient and robust computation – estimating the unimodal distribution involves solving a simple convex optimization problem – and enables more accurate inferences provided that it holds. Second, the method takes as its input two numbers for each test (an effect size estimate, and corresponding standard error), rather than the one number usually used (*p* value, or *z* score). When available, using two numbers instead of one helps account for variation in measurement precision across tests. It also facilitates estimation of effects, and unlike standard FDR methods our approach provides interval estimates (credible regions) for each effect in addition to measures of significance. To provide a bridge between interval estimates and significance measures we introduce the term “local false sign rate” to refer to the probability of getting the sign of an effect wrong, and argue that it is a superior measure of significance than the local FDR because it is both more generally applicable, and can be more robustly estimated. Our methods are implemented in an R package ashr available from http://github.com/stephens999/ashr.

## Introduction

Since its introduction in 1995 by Benjamini and Hochberg [1], the “False Discovery Rate” (FDR) has quickly established itself as a key concept in modern statistics, and the primary tool by which most practitioners handle large-scale multiple testing in which the goal is to identify the non-zero “effects” among a large number of imprecisely-measured effects.

Here we consider an Empirical Bayes (EB) approach to FDR. This idea is, of course, far from new: indeed, the notion that EB approaches could be helpful in handling multiple comparisons predates introduction of the FDR (e.g. [2]). More recently, EB approaches to the FDR have been extensively studied by several authors, especially B. Efron and co-authors [3–7]; see also [8–11] for example.

So what is the “New Deal” here? We introduce two simple ideas that are new (at least compared with existing widely-used FDR pipelines) and can substantially affect inference. The first idea is to *assume that the distribution of effects is unimodal*. We provide a simple, fast, and stable computer implementation for perfoming EB inference under this assumption, and illustrate how it can improve inferences when the unimodal assumption is correct. The second idea is to use two numbers – effect sizes, and their standard errors – rather than just one – *p* values, or *z* scores – to summarize each measurement. Here we use this idea to allow variation in measurement precision to be better accounted for, avoiding a problem with standard pipelines that poor-precision measurements can inflate estimated FDR. ([12] also suggest using more than one number in FDR analyses, taking a rather different approach to the one we use here.)

In addition to these two new ideas, we highlight a third idea that is old, but which remains under-used in practice: the idea that it may be preferable to focus on estimation rather than on testing. In principle, Bayesian approaches can naturally unify testing and estimation into a single framework – testing is simply estimation with some positive prior probability that the effect is exactly zero. However, despite ongoing interest in this area from both frequentist [13] and Bayesian [14,15] perspectives, in practice large-scale studies that assess many effects almost invariably focus on testing significance and controlling the FDR, and not on estimation. To help provide a bridge between FDR and estimation we introduce the term “local false sign rate” (*lfsr*), which is analogous to the “local false discovery rate” (*lfdr*) [6], but which measures confidence in the *sign* of each effect rather than confidence in each effect being non-zero. We show that in some settings, particularly those with many discoveries, the *lfsr* and *lfdr* can be quite different, and emphasize benefits of the *lfsr*, particularly its increased robustness to modeling assumptions.

Although we focus here on FDR applications, the idea of performing EB inference using a flexible unimodal prior distribution is useful more generally. For example, the methods described here can be applied directly to perform shrinkage estimation for wavelet denoising [16], an idea explored in a companion paper [17]. And analogous ideas can be used to perform EB inference for variances [18]. Importantly, and perhaps surprisingly, our work demonstrates how EB inference under a general unimodal assumption is, if anything, computationally simpler than commonly-used more restrictive assumptions – such as a spike and slab or Laplace prior distribution [19] – as well as being more flexible.

We refer to our EB method as adaptive shrinkage, or ash, to emphasise its key points: using a unimodal prior naturally results in shrinkage estimation, and the shrinkage is adaptive to both the amount of signal in the data and the measurement precision of each observation. We provide implementations in an R package, ashr, available at http://github.com/stephens999/ashr. Code and instructions for reproducing analyses and figures in this paper are at https://github.com/stephenslab/ash.

## Methods

### Model Outine

Here we describe the simplest version of the method, and briefly discuss embellishments we have also implemented.

Let *β* = (*β*_1_,…, *β_J_*) denote *J* “effects” of interest. For example, in a genomics application *β_j_* might be the difference in the mean (log) expression of gene *j* in two conditions. We tackle both the problem of testing the null hypotheses *H_j_* : *β_j_* = 0, and the more general problem of estimating, and assessing uncertainty in,*β_j_*.

Assume that the available data are estimates 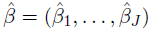 of the effects, and corresponding (estimated) standard errors 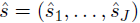. Our goal is to compute a posterior distribution for *β* given 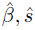, which by Bayes theorem is

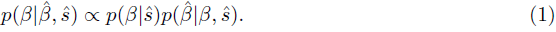

For *p*(*β*|ŝ) we assume that the *β_j_* are independent from a unimodal distribution *g*. This unimodal assumption (UA) is a key assumption that distinguishes our approach from previous EB approaches to FDR analysis. A simple way to implement the UA is to assume that *g* is a mixture of a point mass at 0 and a mixture of *zero-mean* normal distributions:

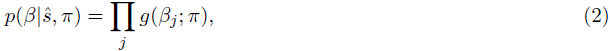

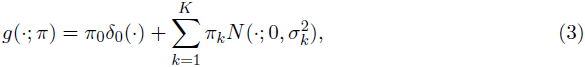

where *δ*_0_(·) denotes a point mass on 0, and N(·; *μ, σ*^2^) denotes the density of the normal distribution with mean *μ* and variance *σ*^2^. Here we take *σ*_1_,…,*σ_K_* to be a large and dense grid of *fixed* positive numbers spanning a range from very small to very big (so *K* is fixed and large). We encourage the reader to think of this grid as becoming infinitely large and dense, as a non-parametric limit, although of course in practice we use a finite grid – see Implementation Details. The mixture proportions π = (π_0_,…, π_*K*_), which are non-negative and sum to one, are hyper-parameters to be estimated.

For the likelihood 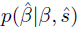 we assume a normal approximation:

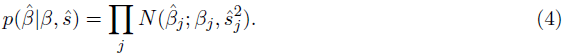

This simple model features both the key ideas we want to emphasize in this paper: the UA is encapsulated in (3) while the different measurement precision of different observations is encapsulated in the likelihood (4) – specifically, observations with larger standard error will have a flatter likelihood, and therefore have less impact on inference. However, the model also has several additional assumptions that can be relaxed. Specifically,

1. The form (3) implies that *g* is symmetric about 0. More flexibility can be obtained using mixtures of uniforms [Equation (12)]; indeed this allows *g* to approximate *any* unimodal distribution.
2. The model (2) assumes that the effects are identically distributed, independent of their standard errors *ŝ*. This can be relaxed [see (11)].
3. The normal likelihood (4) can be generalized to a *t* likelihood [see (13)].

These embellishments, detailed in Detailed Methods, are implemented in our software. Other limitations are harder to relax, most notably the independence and conditional independence assumptions (which are also made by most existing approaches). Correlations among tests certainly arise in practice, either due to genuine correlations or due to unmeasured confounders, and their potential impact on estimated FDRs is important to consider whatever analysis methods are used [20,21].

### Fitting the Model

Together, (2)–(4) imply that 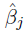 are independent with 
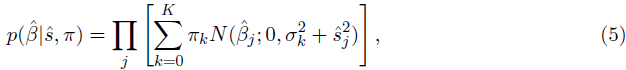

where we define *σ_0_* : = 0.

The usual EB approach to fitting this model would involve two steps:

1. Estimate the hyper-parameters π by maximizing the likelihood *L*(π), given by (5), yielding 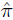 : = argmax *L*(π).
2. Compute quantities of interest from the conditional distributions 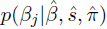. For example, the evidence against the null hypothesis *β_j_* = 0 can be summarized by 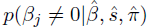.

Both steps are straightforward. Step 1 is a convex optimization problem, and can be solved quickly and reliably using interior point methods [22,23]. (Alternatively a simple EM algorithm can also work well, particularly for modest *J*; see http:://stephenslab.github.io/ash/analysis/IPvsEM.html.) And the conditional distributions 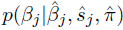 in Step 2 are analytically available, each a mixture of a point mass on zero and *K* normal distributions. The simplicity of step 1 is due to our use of a fixed grid for *σ_k_* in (3), instead of estimating *σ_k_*. This simple device may be useful in other applications.

Here we slightly modify this usual procedure: instead of obtaining 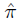 by maximizing the likelihood, we maximize a penalized likelihood [see (18)], where the penalty encourages 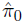 to be as big as possible whilst remaining consistent with the observed data. We introduce this penalty because in FDR applications it is considered desirable to avoid underestimating π_0_ so as to avoid underestimating the FDR.

Our R package implementation typically takes 20s on a modern laptop for *J* = 100, 000, and scales linearly with *J*.

### The Local False Discovery Rate and Local False Sign Rate

As noted above, the posterior distributions 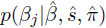 have a simple analytic form. In practice it is common, and desirable, to summarize these distributions to convey the “significance” of each observation *j*. One natural measure of the significance of observation *j* is its “local FDR” [6]

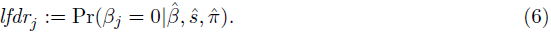

In words, *Ifdr_j_* is the probability, given the observed data, that effect *j* would be a false discovery, if we were to declare it a discovery.

The *Ifdr*, like most measures of significance, is rooted in the hypothesis testing paradigm which focuses on whether or not an effect is exactly zero. This paradigm is popular, despite the fact that many statistical practitioners have argued that it is often inappropriate because the null hypothesis *H_j_* : *β_j_*= 0 is often implausible. For example, Tukey [24] argued that “All we know about the world teaches us that the effects of *A* and *B* are always different – in some decimal place – for any *A* and *B*. Thus asking ‘Are the effects different?’ is foolish.” Instead, Tukey ( [25], p32) suggested that one should address

> …the more meaningful question: “is the evidence strong enough to support a belief that the observed difference has the correct sign?”

Along the same lines, Gelman and co-authors [15,26] suggest focussing on “type S errors”, meaning errors in sign, rather than the more traditional type I errors.

Motivated by these suggestions, we define the “local False Sign Rate” for effect *j, Ifsr_j_*, to be the probability that we would make an error in the sign of effect *j* if we were forced to declare it either positive or negative. Specifically,

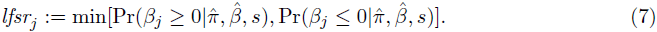

To illustrate, suppose that

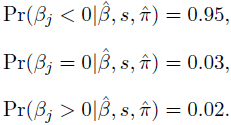

Then from (7) *lfsr_j_* = min(0.05, 0.98) = 0.05 (and, from (6), *lfd_j_* = 0.03). This *lfsr* corresponds to the fact that, given these results, we would guess that *β_j_* is negative, with probability 0.05 of being wrong.

As our notation suggests, *Ifsr_j_* is analogous to *Ifdr_j_*: whereas small values of *Ifdr_j_* indicate that we can be *confident that *β_j_* is non-zero*, small values of *Ifsr_j_* indicate that we can be *confident in the sign of β_j_*. Of course, being confident in the sign of an effect logically implies that we are confident it is non-zero, and this is reflected in the fact that *lfsr_j_ ≥ lfdr_j_* (this follows from the definition because both the events *β_j_* ≥ 0 and *β_j_* ≤ 0 in (7) include the event *β_j_* = 0). In this sense *lfsr* is a more conservative measure of significance than *lfdr*. More importantly, *lfsr* is more robust to modeling assumptions (see Results).

From these “local” measures of significance, we can also compute average error rates over subsets of observations Г ⊂ {1,…, *J*}. For example,

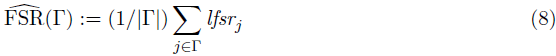

is an estimate of the total proportion of errors made if we were to estimate the sign of all effects in Г (an error measure analogous to the usual (tail) FDR). And, we can define the “*s*-value” 
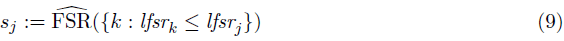

as a measure of significance analogous to Storey's *q*-value [27]. (Replacing *Ifsr_j_* with *Ifdr_j_* in (8) and (9) gives estimates for the usual FDR(Г), and the *q*-values respectively.)

### Related Work

#### Previous Approaches Focussed on FDR

Among previous methods that explicitly consider FDR, our work seems naturally compared with the EB methods of [6] and [11] (implemented in the R packages `locfdr` and `mixfdr` respectively) and with the widely-used methods from [27] (implemented in the R package `qvalue`), which although not formally an EB approach, shares some elements in common.

There are two key differences between our approach and these existing methods. First, whereas these existing methods take as input a single number – either a *z* score (locfdr and mixfdr), or a *p* value (qvalue) – for each effect, we instead work with two numbers (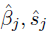). Here we are building on [28], who develops Bayes Factors testing individual null hypotheses in a similar way; see also [29]. Using two numbers instead of one clearly has the potential to be more informative, as illustrated in Results (Figure 4).

Second, our unimodal assumption (UA) that the effects are unimodal about zero, is an assumption not made by existing methods, and one that we argue to be both plausible and beneficial in many contexts. Although the UA will not hold in all settings, it will often be reasonable, especially in FDR-related contexts that focus on rejecting the null hypotheses *β_j_* = 0. This is because if “*β_j_* = 0” is a plausible null hypothesis then “*β_j_* very near 0” should also be plausible. Further, it seems reasonable to expect that larger effects become decreasingly plausible, and so the distribution of the effects will be unimodal about 0. To paraphrase Tukey, “All we know about the world teaches us that large effects are rare, whereas small effects abound.” We emphasize that the UA relates to the distribution of *all* effects, and not only the *detectable* effects (i.e. those that are significantly different from zero). It is very likely that the distribution of *detectable* non-zero effects will be multimodal, with one mode for detectable positive effects and another for detectable negative effects, and the UA does not contradict this.

In further support of the UA, note that large-scale regression methods almost always make the UA for the regression coefficients, which are analogous to the “effects” we estimate here. Common choices of unimodal distribution for regression coefficients include the spike and slab, Laplace, *t*, normal-gamma, normal-inverse-gamma, or horseshoe priors [30]. These are all less flexible than our approach, which provides for general unimodal distributions, and it may be fruitful to apply our methods to the regression context; indeed see [31] for work in this vein. Additionally, the UA can be motivated by its effect on point estimates, which is to “shrink” the estimates towards the mode – such shrinkage is desirable from several standpoints for improving estimation accuracy. Indeed most model-based approaches to shrinkage make parametric assumptions that obey the UA (e.g. [19]).

Besides its plausibility, the UA has two important practical benefits: it facilitates more accurate estimates of FDR-related quantities, and it yields simple algorithms that are both computationally and statistically stable (see Results).

#### Other Work

There is also a very considerable literature that does not directly focus on the FDR problem, but which involves similar ideas and methods. Among these, a paper about deconvolution [32] is most similar, methodologically, to our work here: indeed, this paper includes all the elements of our approach outlined above, except for the point mass on 0 and corresponding penalty term. However, the focus is very different: [32] focuses entirely on estimating *g*, whereas our primary focus is on estimating *β_j_*. Also, they provide no software implementation.

More generally, the related literature is too large to review comprehensively, but relevant key-words include “empirical Bayes”, “shrinkage”, “deconvolution”, “semi-parametric”, “shape-constrained”, and “heteroskedastic”. Some pointers to recent papers in which other relevant citations can be found include [33–36]. Much of the literature focusses on the homoskedastic case (i.e. *ŝ_j_* all equal) whereas we allow for heteroskedasticity. And much of the recent shrinkage-oriented literature focuses only on point estimation of *β_j_*, whereas for FDR-related applications measures of uncertainty are essential. Several recent papers consider more flexible non-parametric assumptions on *g* than the UA assumption we make here. In particular, [35,37] consider the unconstrained non-parametric maximum likelihood estimate (NPMLE) for *g*. These methods may be useful in settings where the UA assumption is considered too restrictive. However, the NPMLE for *g* is a discrete distribution, which will induce a discrete posterior distribution on *β_j_*, and so although the NPMLE may perform well for point estimation, it may not adequately reflect uncertainty in *β_j_*. To address this some regularization on *g* (e.g. as in [36]) may be necessary. Indeed, one way of thinking about the UA is as a way to regularize *g*.

## Results

We compare results of ash with existing FDR-based methods implemented in the R packages `qvalue` (v2.1.1), `locfdr` (v1.1-8), and `mixfdr` (v1.0, from https://cran.r-project.org/src/contrib/Archive/mixfdr/). In all our simulations we assume that the test statistics follow the expected theoretical distribution under the null, and we indicate this to `locfdr` using `nulltype=0` and to `mixfdr` using `theonull=TRUE`. Otherwise all packages were used with default options.

### Effects of the Unimodal Assumption

Here we consider the effects of making the UA. To isolate these effects we consider the simplest case, where every observation has the same standard error, *s_j_* = 1. That is, 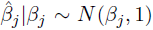 and *ŝ_j_* = *sj* = 1. In this case the *z* scores 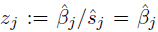, so modelling the *z* scores is the same as modelling the 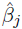. Thus the primary difference among methods in this setting is that ash makes the UA and other methods do not.

To briefly summarize the results in this section:

1. The UA can produce quite different results from existing methods.
2. The UA can yield conservative estimates of the proportion of true nulls, π_0_, and hence conservative estimates of *lfdr* and FDR.
3. The UA yields a procedure that is numerically and statistically stable, and is somewhat robust to deviations from unimodality.

#### The UA can Produce Quite Different Results from Existing Methods

We illustrate the effects of the UA with a simple simulation, with effects *β_j_* ~ *N*(0,1) (so with *s_j_* = 1, 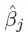 ~ N(0,2)). Though no effects are null, there are many *p* values near 1 and *z* scores near 0 (Figure 1). We used qvalue, locfdr, mixfdr and ash to decompose the *z* scores (*z_j_* = 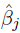), or their corresponding *p* values, into null and alternative components. Here we are using the fact that these methods all provide an estimated *lfdr_j_* for each observation *j*, which implies such a decomposition; specifically the average *lfdr_j_* within each histogram bin estimates the fraction of observations in that bin that come from the null vs the alternative component. The results (Figure 1) illustrate a clear difference between ash and existing methods: the existing methods have a “hole” in the alternative z score distribution near 0, whereas ash, due to the UA, has a mode near 0. (Of course the null distribution also has a peak at 0, and the *lfdr* under the UA is still smallest for *z* scores that are far from zero – i.e. large *z* scores remain the “most significant”.)

**Figure 1.**
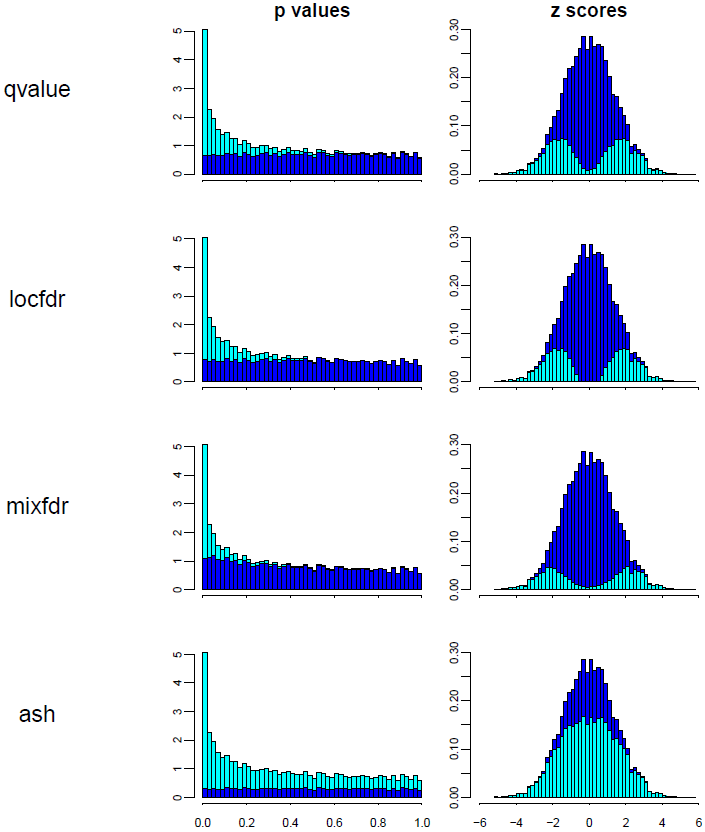
Illustration that the unimodal assumption (UA) in ash can produce very different results from existing methods. The figure shows, for a single simulated dataset, the way different methods decompose *p* values (left) and *z* scores (right) into a null component (dark blue) and an alternative component (cyan). In the *z* score space the alternative distribution is placed on the bottom to highlight the differences in its shape among methods. The three existing methods (`qvalue`, `locfdr`, `mixfdr`) all produce a “hole” in the alternative *z* score distribution around 0. In contrast `ash` makes the UA – that the effect sizes, and thus the *z* scores, have a unimodal distribution about 0 – which yields a very different decomposition. (In this case the ash decomposition is closer to the truth: the data were simulated under a model where all of the effects are non-zero, so the “true” decomposition would make everything cyan.)

This qualitative difference among methods is quite general, and also occurs in simulations where most effects are null (e.g. http://stephenslab.github.io/ash/analysis/referee_uaza.html). To understand why the alternative distribution of *z* scores from locfdr and qvalue has a hole at zero, note that neither of these methods explicitly models the alternative distribution: instead they simply subtract a null distribution (of *z* scores or *p* values) from the observed empirical distribution, letting the alternative distribution be defined implicitly, by what remains. In deciding how much null distribution to subtract – that is, in estimating the null proportion, π_0_ – both methods assume that all *z* scores near zero (or, equivalently, all *p* values near 1) are null. The consequence of this is that their (implicitly-defined) distribution for the alternative *z* scores has a “hole” at 0 – quite different from our assumption of a mode at zero. (Why mixfdr exhibits similar behaviour is less clear, since it does explicitly model the alternative distribution; however we believe it may be due to the default choice of penalty term β described in [11].)

Figure 1 is also helpful in understanding the interacting role of the UA and the penalty term (18) that attempts to make π_0_ as “large as possible” while remaining consistent with the UA. Specifically, consider the panel that shows ash's decomposition of *z* scores, and imagine increasing π_0_ further. This would increase the null component (dark blue) at the expense of the alternative component (light blue). Because the null component is *N*(0,1), and so is biggest at 0, this would eventually create a “dip” in the light-blue histogram at 0. The role of the penalty term is to push the dark blue component as far as possible, right up to (or, to be conservative, just past) the point where this dip appears. In contrast the existing methods effectively push the dark blue component until the light-blue component *disappears* at 0. See https://stephens999.shinyapps.io/unimodal/unimodal.Rmd for an interactive demonstration.

#### *The UA can Produce Conservative Estimates of* π_0_

Figure 1 suggests that the UA will produce smaller estimates of π_0_ than existing methods. Consequently ash will estimate smaller *lfdr*s and FDRs, and so identify more significant discoveries at a given threshold. This is desirable, provided that these estimates remain conservative: that is, provided that 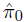 does not underestimate the true π_0_ and 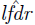 does not underestimate the true *lfdr*. The penalty term (18) aims to ensure this conservative behavior. To check its effectiveness we performed simulations under various alternative scenarios (i.e. various distributions for the non-zero effects, which we denote *g1*), and values for π_0_. The alternative distributions are shown in Figure 2a, with details in Table 2. They range from a “spiky” distribution – where many non-zero β are too close to zero to be reliably detected, making reliable estimation of π_0_ essentially impossible – to a much flatter distribution, which is a normal distribution with large variance (“big-normal”) – where most non-zero β are easily detected making reliable estimation of π_0_ easier. We also include one asymmetric distribution (“skew”), and one clearly bimodal distribution (“bimodal”), which, although we view as generally unrealistic, we include to assess robustness of ash to deviations from the UA.

**Figure 2.**
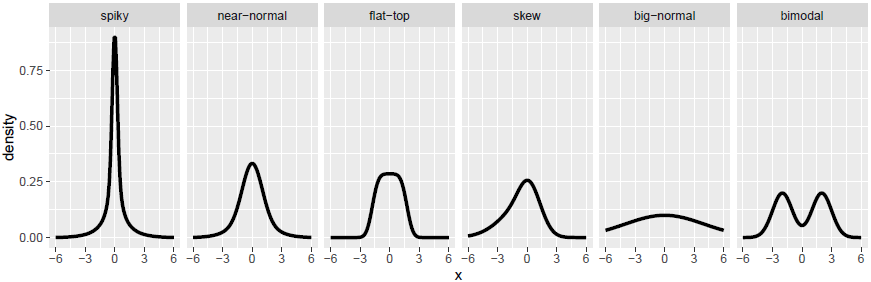
Results of simulation studies (constant precision *s_j_* = 1) (a) Densities of non-zero effects, *g*_1_, used in simulations.

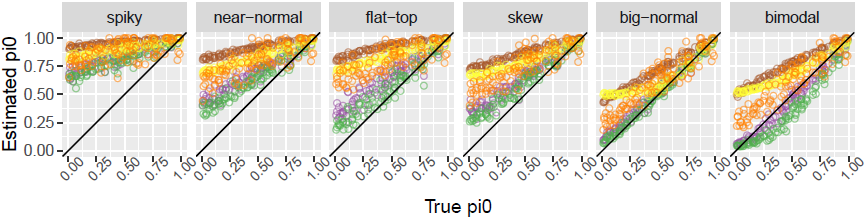
(b) Comparison of true and estimated values of π_0_. When the UA holds all methods typically yield conservative (over-)estimates for π_0_, with ash being least conservative, and hence most accurate. qvalue is sometimes anti-conservative when π_0_ ≈ 1. When the UA does not hold (“bimodal” scenario)
the ash estimates are slightly anti-conservative.

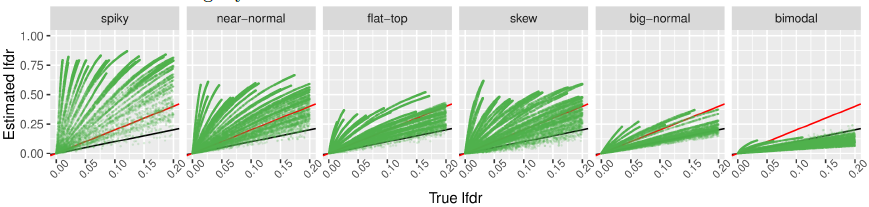
(c) Comparison of true and estimated *lfdr* from ash (ash.n). Black line is *y* = *x* and red line is *y* = 2*x*. Estimates of lfdr are conservative when UA holds, due to conservative estimates of π_0_.

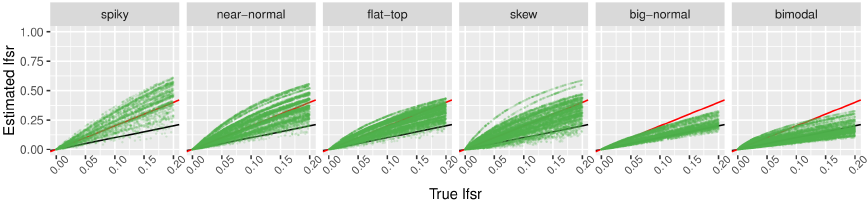
(d) As in c), but for *lfsr* instead of *lfdr*. Estimates of *lfsr* are consistently less conservative than lfdr when UA holds, and also less anti-conservative in bimodal scenario.

**Table 1.** Table of empirical coverage for nominal 95% lower credible bounds. (a) All observations. Coverage rates are generally satisfactory, except for the extreme \spiky" scenario. This is due to the penalty term (18) which tends to cause over-shrinking towards zero. Removing this penalty term produces coverage rates closer to the nominal levels for uniform and normal methods (Table 3). Removing the penalty in the half-uniform case is not recommended (see Appendix).

|  | spiky | near-normal | flat-top | skew | big-normal | bimodal |
| --- | --- | --- | --- | --- | --- | --- |
| ash.n | 0.90 | 0.94 | 0.95 | 0.94 | 0.96 | 0.96 |
| ash.u | 0.87 | 0.93 | 0.94 | 0.93 | 0.96 | 0.96 |
| ash.hu | 0.88 | 0.93 | 0.94 | 0.94 | 0.96 | 0.96 |

(b) “Significant” negative discoveries. Coverage rates are generally satisfactory, except for the uniform-based methods in the spiky and near-normal scenarios, and the normal-based method in the at-top scenario. These results likely reect inaccurate estimates of the tails of *g* due to a disconnect between the tail of *g* and the component distributions in these cases. For example, the uniform methods sometimes substantially underestimate the length of the tail of *g* in these long-tailed scenarios, causing over-shrinkage of the tail toward 0.

|  | spiky | near-normal | flat-top | skew | big-normal | bimodal |
| --- | --- | --- | --- | --- | --- | --- |
| ash.n | 0.93 | 0.94 | 1.00 | 0.94 | 0.95 | 0.98 |
| ash.u | 0.88 | 0.90 | 0.93 | 0.91 | 0.94 | 0.94 |
| ash.hu | 0.87 | 0.87 | 0.92 | 0.93 | 0.94 | 0.94 |

“Significant” positive discoveries. Coverage rates are generally satisfactory, except for the symmetric methods under the asymmetric (“skew”) scenario.

|  | spiky | near-normal | flat-top | skew | big-normal | bimodal |
| --- | --- | --- | --- | --- | --- | --- |
| ash.n | 0.94 | 0.94 | 0.94 | 0.86 | 0.95 | 0.96 |
| ash.u | 0.93 | 0.93 | 0.93 | 0.84 | 0.95 | 0.95 |
| ash.hu | 0.92 | 0.92 | 0.93 | 0.92 | 0.95 | 0.95 |

For each simulation scenario we simulated 100 independent data sets, each with *J* = 1000 observations. For each data set we simulated data as follows:

1. Simulate π_0_ ~ *U*[0,1].
2. For *j* = 1,…, *J*, simulate *β_j_* ~ π_0_δ_0_ + (1 − π_0_)*g*_1_(·).
3. For *j* = 1,…, *J*, simulate 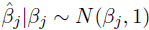.

Figure 2b compares estimates of π_0_ from qvalue, locfdr, mixfdr and ash (*y* axis) with the true values (*x* axis). For ash we show results for *g*1 modelled as a mixture of normal components (“ash.n”) and as a mixture of symmetric uniform components (“ash.u”). (Results using the asymmetric uniforms, which we refer to as “half-uniforms”, and denote “ash.hu” in subsequent sections, are here generally similar to ash.u and omitted to avoid over-cluttering figures.) The results show that ash provides the smallest more accurate, estimates for π_0_, while remaining conservative in all scenarios where the UA holds. When the UA does not hold (“bimodal” scenario) the ash estimates can be slightly anti-conservative. We view this as a minor concern in practice, since we view such a strong bimodal scenario as unlikely in most applications where FDR methods are used. (In addition, the effects on *lfsr* estimates turn out to be relatively modest; see below).

#### The lfsr is More Robust than lfdr

The results above show that ash can improve on existing methods in producing smaller, more accurate, estimates of π_0_, which will lead to more accurate estimates of FDR. Nonetheless, in many scenarios ash continues to substantially over-estimate π_0_ (see the “spiky” scenario for example). This is because these scenarios include an appreciable fraction of “small nonnull effects” that are essentially indistinguishable from 0, making accurate estimation of π_0_ impossible. Put another way, and as is well known, π_0_ is not identifiable: the data can effectively provide an upper bound on plausible values of π_0_, but not a lower bound (because the data cannot rule out that everything is non-null, but with minuscule effects). To obtain conservative behavior we must estimate π_0_ by this upper bound, which can be substantially larger than the true value.

Since FDR-related quantities depend quite sensitively on π_0_, the consequence of this overestimation of π_0_ is corresponding overestimation of FDR (and *lfdr*, and *q* values). To illustrate, Figure 2c compares the estimated *lfdr* from ash.n with the true value (computed using Bayes rule from the true *g*1 and π_0_). As predicted, *lfdr* is overestimated, especially in scenarios which involve many non-zero effects that are very near 0 (e.g. the spiky scenario with π_0_ small) where π_0_ can be grossly overestimated. Of course other methods will be similarly affected by this: those that more grossly overestimate π_0_, will more grossly overestimate *lfdr* and FDR/*q*-values.

The key point we want to make here is that estimation of π_0_, and the accompanying identifiability issues, become substantially less troublesome if we use the local false sign rate *lfsr* (7), rather than *lfdr,* to measure significance. This is because *lfsr* is less sensitive to the estimate of π_0_. To illustrate, Figure 2d compares the estimated *lfsr* from ash.n with the true value: although the estimated *lfsr* continue to be conservative, overestimating the truth, the overestimation is substantially less pronounced than for the *lfdr*, especially for the “spiky” scenario. Further, in the bi-modal scenario, the anti-conservative behavior is less pronounced in *lfsr* than *lfdr*.

Compared with previous debates, this section advances an additional reason for focussing on the sign of the effect, rather than just testing whether it is 0. In previous debates authors have argued against testing whether an effect is 0 because it is *implausible that effects are exactly 0*. Here we add that *even if one believes that some effects may be exactly zero*, it is still better to focus on the sign, because generally *the data are more informative about that question* and so inferences are more robust to, say, the inevitable mis-estimation of π_0_. To provide some intuition, consider an observation with a *z* score of 0. The *lfdr* of this observation can range from 0 (if π_0_ = 0) to 1 (if π_0_ = 1). But, assuming a symmetric *g*, the *lfsr* > 0.5 whatever the value of π_0_, because the observation *z* = 0 says nothing about the sign of the effect. Thus, there are two reasons to use the *lfsr* instead of the *lfdr*: it answers a question that is more generally meaningful (e.g. it applies whether or not zero effects truly exist), and estimation of *lfsr* is more robust.

Given that we argue for using *lfsr* rather than *lfdr*, one might ask whether we even need a point mass on zero in our analysis. Indeed, one advantage of the *lfsr* is that it makes sense even if no effect is exactly zero. And, if we are prepared to assume that no effects are exactly zero, then removing the point mass yields smaller and more accurate estimates of *lfsr* when that assumption is true (Figure 6a). However, there is “no free lunch”: if in fact some effects are exactly zero then the analysis with no point mass will tend to be anti-conservative, underestimating *lfsr* (Figure 6b). We conclude that *if* ensuring a “conservative” analysis is important then one should allow for a point mass at 0.

#### The UA Helps Provide Reliable Estimates of g

An important advantage of our EB approach based on modelling the effects *β_j_*, rather than *p* values or *z* scores, is that it can estimate the effects *β_j_*. Specifically, it provides a posterior distribution for each *β_j_*, which can be used to construct interval estimates, etc. Further, because the posterior distribution is, by definition, conditional on the observed data, interval estimates based on posterior distributions are also valid Bayesian inferences for any subset of the effects that have been selected based on the observed data. This kind of “post-selection” validity is much harder to achieve in the frequentist paradigm. In particular the posterior distribution solves the (Bayesian analogue of the) “False Coverage Rate” problem posed by [13] which [6] summarizes as follows: “having applied FDR methods to select a set of nonnull cases, how can confidence intervals be assigned to the true effect size for each selected case?”. [6] notes the potential for EB approaches to tackle this problem, and [14] considers in detail the case where the non-null effects are normally distributed.

The ability of the EB approach to provide valid “post-selection” interval estimates is extremely attractive in principle. But its usefulness in practice depends on reliably estimating the distribution *g*. Estimating *g* is a “deconvolution problem”, which are notoriously difficult in general. Indeed, Efron emphasizes the difficulties of implementing a stable general algorithm, noting in his rejoinder “the effort foundered on practical difficulties involving the perils of deconvolution … Maybe I am trying to be overly nonparametric … but it is hard to imagine a generally satisfactory parametric formulation …” ([6] rejoinder, p46). We argue here that the UA can greatly simplify the deconvolution problem, producing both computationally and statistically stable estimates of *g*.

To illustrate, we compare the estimated *g* from ash (under the UA) with the non-parametric maximum likelihood estimate (NPMLE) for *g* (i.e. estimated entirely non-parametrically without the unimodal constraint). The NPMLE is straightforward to compute in R using the `REBayes::GLmix` function [35]. Figure 3 shows results under six different scenarios. The estimated cdf from ash is generally closer to the truth, even in the bi-modal scenario. (ash tends to systematically overestimate the mass of *g* near zero; this can be avoided by removing the penalty term (18); Figure 5.) Furthermore, the estimate from ash is also substantially more “regular” than the NPMLE, which has several almost-vertical segments indicative of a concentration of density in the estimated *g* at those locations. Indeed the NPMLE is a discrete distribution [35], so this kind of concentration will always occur. The UA prevents this concentration, effectively regularizing the estimated *g*. While the UA is not the only way to achieve this (e.g. [36]), we view it as attractive and widely applicable.

**Figure 3.**
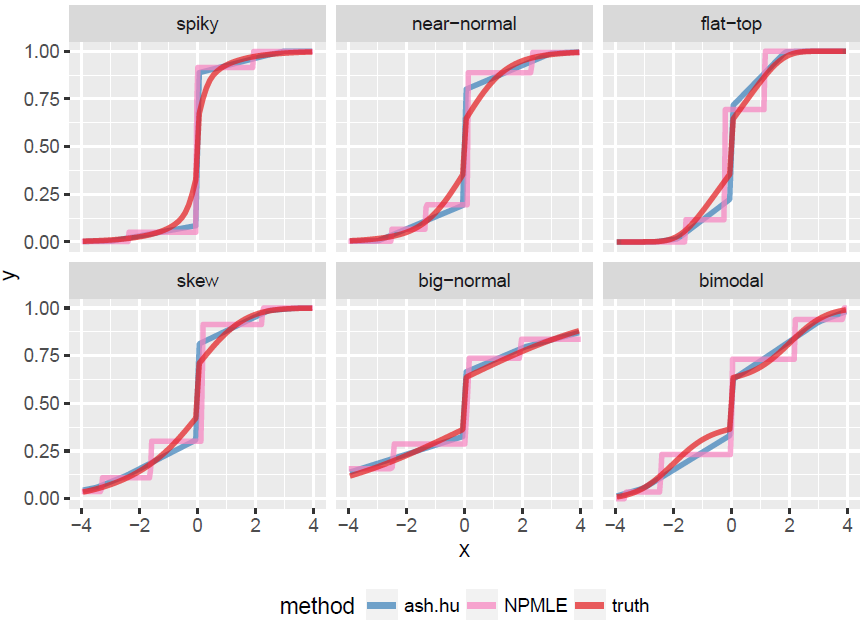
Comparison of estimated cdfs from ash and the NPMLE. Different ash methods perform similarly, so only ash.hu is shown for clarity. Each panel shows results for a single example data set, one for each scenario in Figure 2a. The results illustrate how the unimodal assumption made by ash regularizes the estimated cdfs compared with the NPMLE.

#### Calibration of Posterior Intervals

To quantify the effects of errors in estimates of *g* we examine the calibration of the resulting posterior distributions (averaged over 100 simulations in each Scenario). Specifically we examine the empirical coverage of nominal lower 95% credible bounds for a) all observations; b) significant negative discoveries; c) significant positive discoveries. We examine only lower bounds because the results for upper bounds follow by symmetry (except for the one asymmetric scenario). We separately examine positive and negative discoveries because the lower bound plays a different role in each case: for negative discoveries the lower bound is typically large and negative and limits how big (in absolute value) the effect could be; for positive discoveries the lower bound is positive, and limits how small (in absolute value) the effect could be. Intuitively, the lower bound for negative discoveries depends on the accuracy of *g* in its tail, whereas for positive discoveries it is more dependent on the accuracy of *g* in the center.

The results are shown in Table 1. Most of the empirical coverage rates are in the range 0.92-0.96 for nominal coverage of 0.95, which we view as adequate for practical applications. The strongest deviations from nominal rates are noted and discussed in the table captions. One general issue is that the methods based on mixtures of uniform distributions often slightly curtail the tail of *g*, causing the probability of very large outlying effects to be understated; see also http://stephenslab.github.io/ash/analysis/efron.fcr.html.

### Differing Measurement Precision Across Units

We turn now to the second important component of our work: allowing for varying measurement precision across units. The key to this is the use of a likelihood, (4) or (13), that explicitly incorporates the measurement precision (standard error) of each 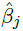.

To illustrate, we conduct a simulation where half the measurements are quite precise (stan-dard error *s_j_* = 1), and the other half are very poor (*s_j_* = 10). In both cases, we assume that half the effects are null and the other half are normally distributed with standard deviation 1:

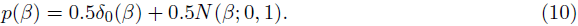

In this setting, the poor-precision measurements (*s_j_* = 10) tell us very little, and any sane analysis should effectively ignore them. However, this is not the case in standard FDR-type analyses (Figure 4). This is because the poor measurements produce *p* values that are approximately uniform (Figure 4a), which, when combined with the good-precision measurements, dilute the overall signal (e.g. they reduce the density of *p* values near 0). This is reflected in the results of FDR methods like `qvalue` and `locfdr`: the estimated error rates (*q*-values, or *Ifdr* values) for the good-precision observations increase when the low-precision observations are included in the analysis (Figure 4b). In contrast, the results from ash for the good-precision observations are unaffected by including the low-precision observations in the analysis (Figure 4b).

**Figure 4.**
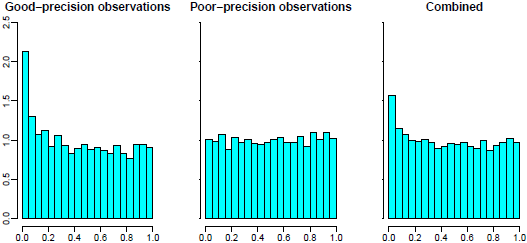
Simulations showing how, with existing methods, but not ash, poor-precision observations can contaminate signal from good-precision observations. (a) Density histograms of *p* values for good-precision, poor-precision, and combined observations. The combined data show less signal than the good-precision data, due to the contamination effect of the poor-precision measurements.

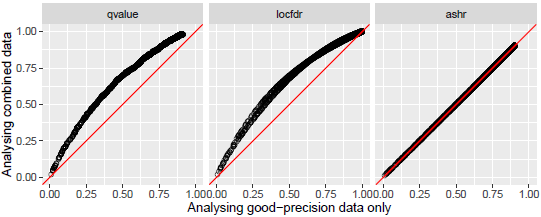
(b) Results of different methods applied to good-precision observations only (*x* axis) and combined data (*y* axis). Each point shows the “significance” (*q* values from qvalue; *lfdr* for locfdr; *lfsr* for ash) of a good-precision observation under the two different analyses. For existing methods including the poor-precision observations reduces significance of good-precision observations, whereas for ash the poor-precision observations have little effect (because they have a very at likelihood).

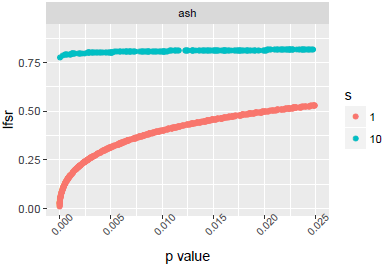
(c) The relationship between lfsr and p-value is different for good-precision (*s* = 1) and low-precision (*s* = 10) measurements: ash assigns the low-precision measurements a higher lfsr, effectively downweighting them.

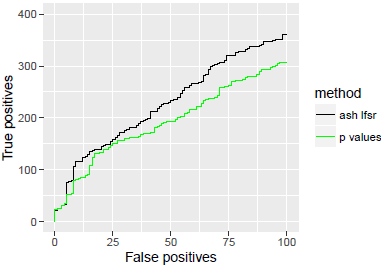
(d) Trade-off between true positives (*y*) vs false positives (*x*) as the significance threshold (lfsr or *p* value) is varied. By downweighting the low-precision observations ash re-orders the significance of observations, producing more true positives at a given number of false positives.

Another consequence of incorporating measurement precision into the likelihood is that ash can re-order the significance of observations compared with the original *p* values or *z* scores. Effectively ash downweights the poor-precision observations by assigning them a higher *Ifsr* than good precision measurements that have the same *p* value (Figure 4c). The intuition is that, due to their poor precision, these measurements contain very little information about the sign of the effects (or indeed any other aspect of the effects), and so the *Ifsr* for these poor-precision measurements is always high; see [38] for related discussion. Here this re-ordering results in improved performance: ash identifies more true positive effects at a given level of false positives (Figure 4d).

#### Dependence of β_j_ on s_j_

Although downweighting low-precision observations may seem intuitive, we must now confess that the issues are more subtle than our treatment above suggests. Specifically, it turns out that the downweighting behaviour depends on an assumption that we have made up to now, that */ j* is independent of *s_j_* (Equation (2)). In practice this assumption may not hold. For example, in gene expression studies, genes with higher biological variance may tend to exhibit larger effects *β_j_* (because they are less constrained). These genes will also tend to have larger *s_j_*, inducing a dependence between *β_j_* and *s_j_*.

Motivated by this, we generalize the prior (2) to 
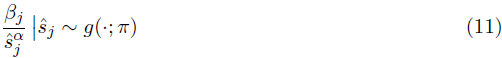

where α is to be estimated or specified. Setting α = 0 yields (2). Setting α > 0 implies that the effects with larger standard error tend to be larger (in absolute value). Fitting this model for any *α* is straightforward using the same methods as for *α* = 0 (see Methods).

The case *α* = 1 in (11) is of special interest because it corresponds most closely to existing methods. Specifically, it can be shown that with *α* = 1 the *lfs_j_* values from ash.n are monotonic in the *p* values: effects with smaller *p* values have smaller *lfsr_j_*. This result generalizes a result in [28], who referred to a prior that produces the same ranking as the *p* values as a “*p* value prior”. Of course, if the *lfsr_j_* are monotonic in the *p* values then the downweighting and reordering of significance illustrated in Figure 4 will not occur. The intuition is that under *α* = 1 the poor precision observations have larger effects sizes, and consequently the same power as the high-precision observations – under these conditions the poor precision observations are not “contaminating” the high precision observations, and so downweighting them is unnecessary. Thus running ash with *α* = 1 will produce the same significance ranking as existing methods. Nonetheless, it is not equivalent to them, and indeed still has the benefits outlined previously: due to the UA ash can produce less conservative estimates of *π_0_* and *lfdr_j_*; and because ash models the *β_j_* it can produce interval estimates.

As an aside, we note that with *α* = 1 the ash estimates of both *g* and *lfsr_j_* depend on the pairs (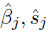) only through the *z* scores 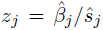 (though interval estimates for *β_j_* still depend on (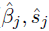)). This means that ash can be run (with *α* = 1) in settings where the *z_j_* are available but the (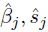) are not. It also opens the intriguing possibility of running ash on *p* values obtained from any test (e.g. a permutation test), by first converting each *p* value to a corresponding *z* score. However, the meaning of and motivation for the UA may be unclear in such settings, and caution seems warranted before proceeding along these lines.

In practice, appropriate choice of *α* will depend on the actual relationship between *β_j_* and *s_j_*, which will be dataset-specific. Further, although we have focussed on the special cases *α* = 0 and *α* = 1, there seems no strong reason to expect that either will necessarily be optimal in practice. Following the logic of the EB approach we suggest selecting *α* by maximum likelihood, which is implemented in our software using a simple 1-d grid search (implementation due to C. Dai).

## Discussion

We have presented an Empirical Bayes approach to large-scale multiple testing that emphasizes two ideas. First, we emphasize the potential benefits of using two numbers (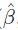, and its standard error) rather than just one number (a *p* value or *z* score) to summarize the information on each test. While requiring two numbers is slightly more onorous than requiring one, in many settings these numbers are easily available and if so we argue it makes sense to use them. Second, we note the potential benefits – both statistical and computational – of assuming that the effects come from a unimodal distribution, and provide flexible implementations for performing inference under this assumption. We also introduce the “false sign rate” as an alternative measure of error to the FDR, and illustrate its improved robustness to errors in model fit, particularly mis-estimation of the proportion of null tests, π_0_.

Multiple testing is often referred to as a “problem” or a “burden”. In our opinion, EB approaches turn this idea on its head, treating multiple testing as an *opportunity*: specifically, an opportunity to learn about the prior distributions, and other modelling assumptions, to improve inference and make informed decisions about significance (see also [2]). This view also emphasizes that, what matters in multiple testing settings is *not* the number of tests, but the *results* of the tests. Indeed, the FDR at a given fixed threshold does not depend on the number of tests: as the number of tests increases, both the true positives and false positives increase linearly, and the FDR remains the same. (If this intuitive argument does not convince, see [27], and note that the FDR at a given *p* value threshold does not depend on the number of tests *m*.) Conversely, the FDR *does* depend on the overall distribution of effects, and particularly on π_0_ for example. The EB approach captures this dependence in an intuitive way: if there are lots of strong signals then we infer π_0_ to be small, and the estimated FDR (or *lfdr*, or *lfsr*) at a given threshold may be low, even if a large number of tests were performed; and conversely if there are no strong signals then we infer π_0_ to be large and the FDR at the same threshold may be high, even if relatively few tests were performed. More generally, overall signal strength is reflected in the estimated *g*, which in turn influences the estimated FDR.

Two important practical issues that we have not addressed here are correlations among tests, and the potential for deviations from the theoretical null distributions of test statistics. These two issues are connected: specifically, unmeasured confounding factors can cause both correlations among tests and deviations from the theoretical null [20,21]. And although there are certainly other factors that could cause dependence among tests, unmeasured confounders are perhaps the most worrisome in practice because they can induce strong correlations among large numbers of tests and profoundly impact results, ultimately resulting in too many hypotheses being rejected and a failure to control FDR. We are acutely aware that, because our method is less conservative than existing methods, it may unwittingly exacerbate these issues if they are not adequately dealt with. Approaches to deal with unmeasured confounders can be largely divided into two types: those that simply attempt to correct for the resulting inflation of test statistics [39,40], and those that attempt to infer confounders using clustering, principal components analysis, or factor models [21,41–43], and then correct for them in computation of the test statistics (in our case, 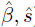). When these latter approaches are viable they provide perhaps the most satisfactory solution, and are certainly a good fit for our framework. Alternatively, our methods could be modified to allow for test statistic inflation, an idea that may be worth pursuing in future work.

Another important practical issue is the challenge of small sample sizes. For example, in genomics applications researchers sometimes attempt to identify differences between two conditions based on only a handful of samples in each. In such settings the normal likelihood approximation (4) will be inadequate. And, although the *t* likelihood (13) partially addresses this issue, it is also, it turns out, not entirely satisfactory. The root of the problem is that, with small sample sizes, raw estimated standard errors *ŝ_j_* can be horribly variable. In genomics it is routine to address this issue by applying EB methods [44] to “moderate” (i.e. shrink) variance estimates, before computing *p* values from “moderated” test statistics. We are currently investigating how our methods should incorporate such “moderated” variance estimates to make it applicable to small sample settings.

Our approach involves compromises between flexibility, generality, and simplicity on the one hand, and statistical efficiency and principle on the other. For example, in using an EB approach that uses a point estimate for *g*, rather than a fully Bayes approach that accounts for uncertainty in *g*, we have opted for simplicity over statistical principle. And in summarizing every test by two numbers and making a normal or *t* approximation to the likelihood, we have aimed to produce generic methods that can be applied whenever such summary data are available – just as qvalue can be applied to any set of *p* values for example – although possibly at the expense of statistical efficiency compared with developing multiple tailored approaches based on context-specific likelihoods. Any attempt to produce generic methods will involve compromise between generality and efficiency. In genomics, many analyses – not only FDR-based analyses – involve first computing a series of *p* values before subjecting them to some further downstream analysis. An important message here is that working with two numbers (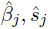), rather than one (*p_j_* or *z_j_*), can yield substantial gains in functionality (e.g. estimating effect sizes, as well as testing; accounting for variations in measurement precision across units) while losing only a little in generality. We hope that our work will encourage development of methods that exploit this idea in other contexts.

## Detailed Methods

### Embellishments

#### More Flexible Unimodal Distributions

Using a mixture of zero-centered normal distributions for *g* in (3) implies that *g* is not only unimodal, but also symmetric. Furthermore, even some symmetric unimodal distributions, such as those with a flat top, cannot be well approximated by a mixture of zero-centered normals.

Therefore, we have implemented a more general approach based on 
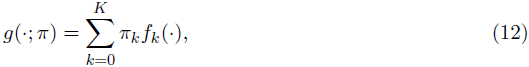

where *f_0_* is a point mass on 0, and *f_k_* (k = *1,…, K*) are pre-specified component distributions with one of the following forms:

1. 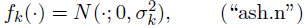
2. 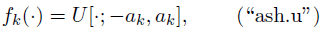
3. 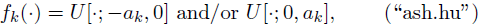
 where *U*[~; *a, b*] denotes the density of a uniform distribution on [*a, b*]. (In (iii) we include both components in the mixture (12), so a grid of values *a_1_,…, a_K_* defines 2*K* + 1 mixture component densities, and π is a 2*K* + 1 vector that sums to 1.) The simplest version (3) corresponds to (i). Replacing these with uniform components (ii)–(iii) only slightly complicates calculations under the normal likelihood (4), and greatly simplifies the calculations under the t likelihood (13) introduced below. The use of uniform components here closely mirrors [32]. (In fact our implementation can handle *any* pre-specified uniform or normal distributions for *f_k_* provided they are all from the same family; however, we restrict our attention here to (i)–(iii) which imply a unimodal *g*.)

Moving from (i) to (iii) the representation (12) becomes increasingly flexible. Indeed, using a large dense grid of 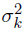 or *a_k_*, (i)–(iii) can respectively approximate, with arbitrary accuracy,

i. any scale mixture of normals, which includes as special cases the double exponential (Laplace) distribution, any *t* distribution, and a very large number of other distributions used in high-dimensional regression settings.
ii. any symmetric unimodal distribution about 0.
iii. any unimodal distribution about 0.

The latter two claims are related to characterizations of unimodal distributions due to [45] and [46]; see [47], p158. In other words, (ii) and (iii) provide fully non-parametric estimation for *g* under the constraints that it is (ii) both unimodal and symmetric, or (iii) unimodal only.

Although our discussion above emphasizes the use of large *K*, in practice modest values of *K* can provide reasonable performance. The key point is that the value of *K* is not critical provided it is sufficiently large, and the grid of *σ_k_* or *a_k_* values suitably chosen. See Implementation for details of our software defaults.

#### Dependence of Effects on Standard Errors

The model (11) for general *α* can be fitted using the algorithm for *α* = 0. To see this, define 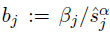, and 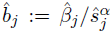. Then 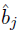 is an estimate of *b_j_* with standard error 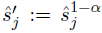. Applying the algorithm for *α* = 0 to effect estimates 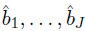, with standard errors 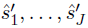 yields a posterior distribution 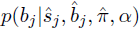, which induces a posterior distribution on 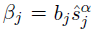.

#### Replace Normal Likelihood with t Likelihood

We generalize the normal likelihood (4) by replacing it with a *t* likelihood: 
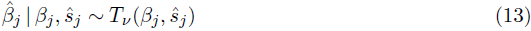

where *T_ν_* (*β_j_, ŝ_j_*) denotes the distribution of *β_j_ + ŝ_j_ T_ν_* where *T_ν_* has a standard *t* distribution on ν degrees of freedom, and ν denotes the degrees of freedom used to estimate *ŝ_j_* (assumed known, and for simplicity assumed to be the same for each *j*). The normal approximation (4) corresponds to the limit ν → ∞. This generalization does not complicate inference when the mixture components *f_k_* in (12) are uniforms; see Implementation below. When the *f_k_* are normal the computations with a *t* likelihood are considerably more difficult and we have not implemented this combination.

Equation (13) is, of course, motivated by the standard asymptotic result

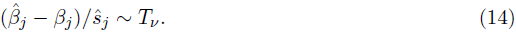

However (14) does not imply (13), because in (14) *ŝ_j_* is random whereas in (13) it is conditioned on. In principle it would be preferable, for a number of reasons, to model the randomness in *ŝ_j_*; we are currently pursuing this improved approach (joint work with M.Lu) and results will be published elsewhere.

#### Non-Zero Mode

An addition to our software implementation, due to C.Dai, allows the mode to be estimated from the data by maximum likelihood, rather than fixed to 0. This involves a simple grid search.

## Implementation Details

### Likelihood for π

We define the likelihood for π to be the probability of the observed data 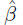 conditional on ŝ: *L*(π) : = 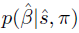, which by our conditional independence assumptions is equal to the product 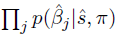. [One might prefer to define the likelihood as 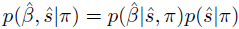, in which case our definition comes down to assuming that the term *p*(ŝ|π) does not depend on π.]

Using the prior 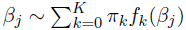 given by (12), and the normal likelihood (4), integrating over *β_j_* yields 
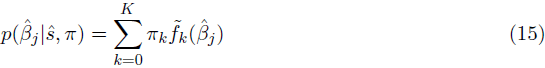

where 
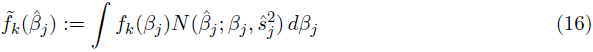

denotes the convolution of *f_k_* with a normal density. These convolutions are straightforward to evaluate whether *f_k_* is a normal or uniform density. Specifically, 
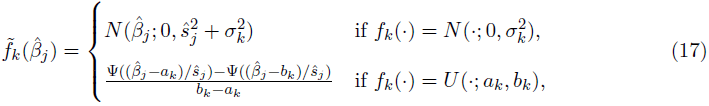

where Ψ denotes the cumulative distribution function (c.d.f.) of the standard normal distribution. If we replace the normal likelihood with the *t_ν_* likelihood (13) then the convolution for f_k_ uniform the convolution is still given by (17) but with Ψ the c.d.f. of the *t_ν_* distribution function. (The convolution for *f_k_* normal is tricky and we have not implemented it.)

### Penalty Term, on π

To make *lfdr* and *lfsr* estimates from our method “conservative” we add a penalty term *log*(*h*(π; λ)) to the log-likelihood log *L*(π) to encourage over-estimation of π_0_: 
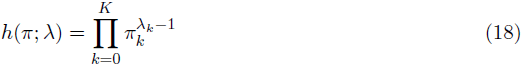

where λ_*k*_ ≥ 1 ∀_*k*_. The default is λ_0_ = 10 and λ_k_ = 1, which yielded consistently conservative estimation of π_0_ in our simulations (Figure 2b).

Although this penalty is based on a Dirichlet density, we do not interpret this as a “prior distribution” for n: we chose it to provide conservative estimates of π_0_ rather than to represent prior belief.

### Problems with Removing the Penalty Term in the Half-Uniform Case

It is straightforward to remove the penalty term by setting λ_*k*_ = 1 in (18). We note here an unanticipated problem we came across when using no penalty term in the half-uniform case (i.e. *f_k_*(·) = *U*[·; — *a_k_*, 0] and/or *U*[·; 0, *a_k_*] in (12)): when the data are nearly null, the estimated *g* converges, as expected and desired, to a distribution where almost all the mass is near 0, but sometimes all this mass is concentrated almost entirely just to one side (left or right) or 0. This can have a very profound effect on the local false sign rate: for example, if all the mass is just to the right of 0 then all observations will be assigned a very high probability of being positive (but very small), and a (misleading) low local false sign rate. For this reason we do not recommend use of the half-uniform with no penalty.

### Optimization

With this in place, the penalized log-likelihood for π is given by: 
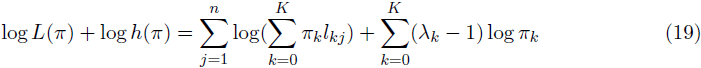

where the 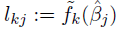 are known. This is a convex optimization problem, which can be solved very quickly and reliably using interior point (IP) methods. We used the KWdual function from the R package REBayes [48], which uses Rmosek [49]. We also found a simple EM algorithm [50], accelerated using the elegant R package SQUAREM [51], to provide adequate performance. In our EM implementation we initialized *π_k_* = *1/n* for *k* = 1,…, *K*, with π_0_ = 1 — π_1_ —…— π_*K*_, and the one-step updates are:

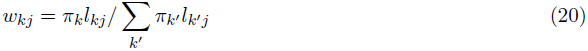

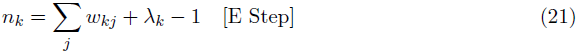

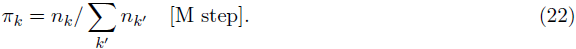

One benefit to the EM algorithm is fewer software dependencies. Both EM and IP methods are implemented in the ashr package; results shown here are from the IP method, but graphs from EM are essentially the same. See http://stephenslab.github.io/ash/analysis/checkIP.html and http://stephenslab.github.io/ash/analysis/IPvsEM.html for comparisons.

### Conditional Distributions

Given 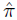, we compute the conditional distributions

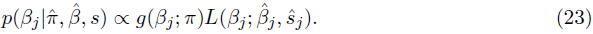

Each posterior is a mixture on *K* + 1 components: 
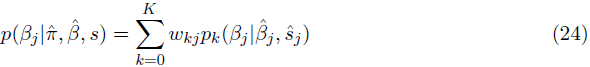

where the posterior weights *w_kj_* are computed as in (20) with π = 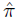, and the posterior mixture component *p_k_* is the posterior on *β_j_* that would be obtained using prior *f_k_* (*β_j_*) and likelihood *L*(*β_j_*; 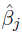, *ŝ_j_*). All these posterior distributions are easily available. For example, if *f_k_* is uniform and *L* is *t_ν_* then this is a truncated t distribution. If *f_k_* is normal and *L* is normal, then this is a normal distribution.

### Choice of Grid for σ_k_, a_k_

When *f_k_* is *N*(0,σ_k_) we specify our grid by specifying: i) a maximum and minimum value (σ_min_,σ_max_); ii) a multiplicative factor *m* to be used in going from one grid-point to the other, so that *σ_k_* = *mσ*_*k−1*_. The multiplicative factor affects the density of the grid; we used *m* = 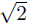 as a default. We chose σ_min_ to be small compared with the measurement precision (σ_min_ = min(*ŝ_j_*)/10) and σ_max_ = 2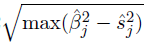 based on the idea that σ_max_ should be big enough so that 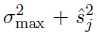 should exceed 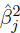. (In rare cases where 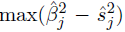 is negative we set σ_max_ = 8σ_min_.)

When the mixture components *f_k_* are uniform, we use the same grid for the parameters *a_k_* as for *σ_k_* described above.

Our goal in specifying a grid was to make the limits sufficiently large and small, and the grid sufficiently dense, that results would not change appreciably with a larger or denser grid. For a specific data set one can of course check this by experimenting with the grid, but these defaults usually work well in our experience.

## Acknowledgements

I thank J. Lafferty for pointing out the convexity of the likelihood function, which lead to an improved implementation with interior point methods. Statistical analyses were conducted in the R programming language [52], Figures produced using the ggplot2 package [53], and text prepared using LTEX. Development of the methods in this paper was greatly enhanced by the use of the knitr package [54] within the RStudio GUI, and git and github. The ashr R package is available from http://github.com/stephens999/ashr and includes contributions from Chaoxing (Rick) Dai, Mengyin Lu, Nan Xiao, and Tian Sen.

This work was supported by NIH grant HG02585 and a grant from the Gordon and Betty Moore Foundation.

## Appendix: Additional Figures and Tables

**Figure 5.**
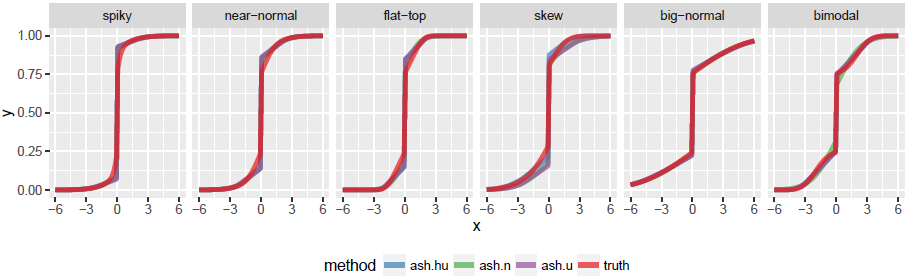
Comparisons of average estimated cdfs of *g* with and without penalty term. See Figure 2b for simulation scenarios. In most cases the three different ash methods are very similar and so the lines lie on top of one another. (a) Average estimated cdfs across ~ 10 data sets compared with truth; methods here use penalty (18) which leads to systematic overestimation of π_0_ in some scenarios.

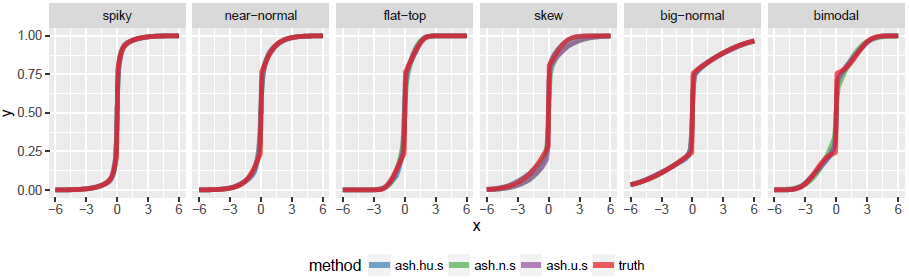
(b) Average estimated cdfs across ~ 10 data sets compared with truth; methods here do not use penalty (18) so π_0_ is not systematically overestimated. Systematic differences from the truth in “skew” and “bimodal” scenarios highlight the effects of model mis-specification.

**Figure 6.**
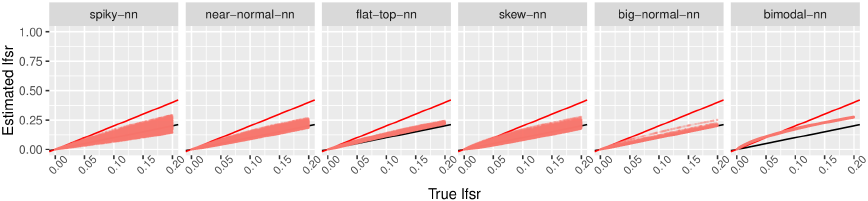
Illustration of effects of excluding a point mass from the analysis. (a) Comparison of true and estimated lfsr when data are simulated with no point mass at zero (π_0_ = 0), and also analyzed by ash with no point mass on 0 (and mixture of normal components for g). Black line is *y* = *x* and red line is *y* = 2*x*. The results illustrate how estimates of lfsr can be more accurate in this case. That is, assuming there is no point mass can be beneficial if that is indeed true.

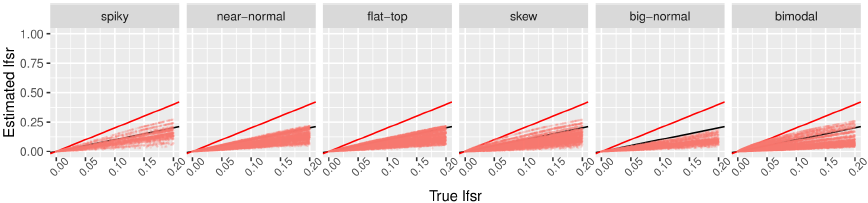
(b) Comparison of true and estimated lfsr when data are simulated with point mass at zero (drawn uniformly from [0,1] in each simulation), but analyzed by ash with no point mass on 0 (and mixture of normal components for *g*). Black line is *y* = *x* and red line is *y* = 2*x*. The results illustrate how estimates of lfsr can be anti-conservative if we assume there is no point mass when the truth is that there is a point mass.

**Table 2.**
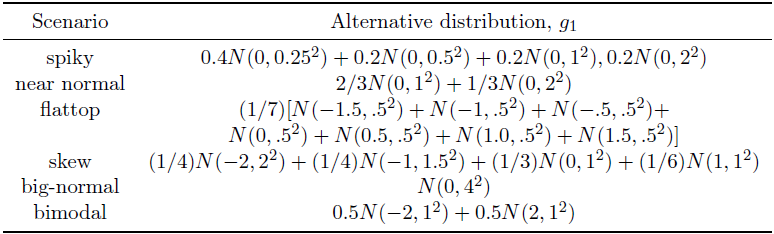
Summary of simulation scenarios considered

| Scenario | Alternative distribution, $g_1$ |
| --- | --- |
| spiky | $0.4N(0, 0.25^2) + 0.2N(0, 0.5^2) + 0.2N(0, 1^2), 0.2N(0, 2^2)$ |
| near normal | $2/3N(0, 1^2) + 1/3N(0, 2^2)$ |
| flattop | $(1/7)[N(-1.5, .5^2) + N(-1, .5^2) + N(-.5, .5^2) + N(0, .5^2) + N(0.5, .5^2) + N(1.0, .5^2) + N(1.5, .5^2)]$ |
| skew | $(1/4)N(-2, 2^2) + (1/4)N(-1, 1.5^2) + (1/3)N(0, 1^2) + (1/6)N(1, 1^2)$ |
| big-normal | $N(0, 4^2)$ |
| bimodal | $0.5N(-2, 1^2) + 0.5N(2, 1^2)$ |

**Table 3.**
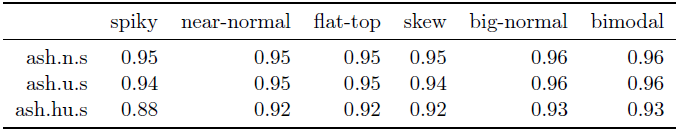
Table of empirical coverage for nominal 95% lower credible bounds for methods *without* the penalty term) (a) All observations

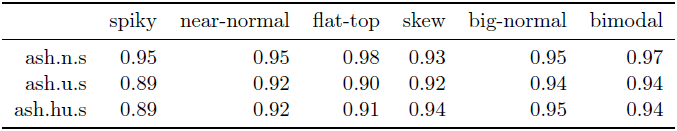
(b) “Significant” negative discoveries.

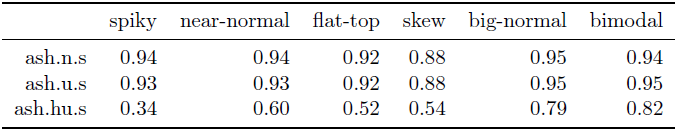
(c) “Significant” positive discoveries.

